# Q-SNARE Syntaxin 7 confers actin-dependent rapidly replenishing synaptic vesicles upon high activity

**DOI:** 10.1101/2020.08.13.249672

**Authors:** Yasunori Mori, Koichiro Takenaka, Yugo Fukazawa, Shigeo Takamori

**Author notes:** All correspondence should be addressed to Y.M. or S.T.

## Abstract

Replenishment of readily releasable synaptic vesicles (SVs) with vesicles in the recycling pool is important for sustained transmitter release during repetitive stimulation. Kinetics of replenishment and available pool size define synaptic performance. However, whether all SVs in the recycling pool are recruited for release with equal probability is unknown. Here, using comprehensive optical imaging for various presynaptic endosomal SNARE proteins in cultured hippocampal neurons, we demonstrate that part of the recycling pool bearing the endosomal Q–SNARE Syntaxin 7 (Stx7) is preferentially mobilized for release during high–frequency repetitive stimulation. Recruitment of the SV pool marked with the Stx7–reporter requires high intra–terminal Ca^2+^ concentrations and actin polymerization. Furthermore, disruption of Stx7 function by overexpressing the N–terminal domain selectively abolished this pool. Thus, our data indicate that endosomal membrane fusion involving Stx7 is essential for adaptation of synapses to respond high-frequency repetitive stimulation.

---

Hundreds to tens of thousands of neurotransmitter–filled synaptic vesicles (SVs), which are morphologically homogeneous, are clustered at presynaptic terminals, potentially capable of releasing their contents upon the arrival of electrical stimuli (Sudhof, 2004). Yet, only a fraction of all SVs is usually available for stimulus–dependent release (Alabi & Tsien, 2012; Rizzoli & Betz, 2005) (but see, Ikeda & Bekkers, 2009). During the last few decades, the three–pool model has emerged, in which neurotransmitter–filled synaptic vesicles are classified into three functionally distinct pools: the readily releasable pool (RRP), the recycling pool that replenishes the RRP during sustained activity, and the reserve pool (RP, also referred to as the resting pool) that rarely participates in recycling (Alabi & Tsien, 2012; Rizzoli & Betz, 2005). Despite intensive research, however, the molecular underpinnings of the three pools have remained elusive.

Exocytic fusion of SVs to the plasma membrane is mediated by formation of so–called SNAP-receptor (SNARE) complexes, consisting of vesicular SNARE synaptobrevin (Syb)/VAMP2 and two target SNAREs Syntaxin 1 (Stx1) and SNAP–25 (Jahn & Scheller, 2006). Considering the extraordinary speed of SV exocytosis in response to action potentials (typically within several ms), RRP is believed to dock physically to the plasma membrane prior to fusion and the canonical neuronal SNAREs described above are responsible for their docked status (Alabi & Tsien, 2012; Rizzoli & Betz, 2005). Aside from the necessity of the dominant SNAREs for the final step of SV exocytosis, non-canonical SNAREs that mediate fusion reactions between intracellular organelles in non–neuronal cells are not only present, but also enriched in SVs purified from rat brains (Takamori *et al*, 2006)(Appendix Fig S1), albeit at lower expression levels than the canonical SNAREs (Wilhelm *et al*, 2014), making them likely candidate protein constituents in functionally distinct subsets of SVs, such as the recycling and reserve pools. Indeed, endosomal SNAREs are sorted to distinct vesicle pools, namely in RP or spontaneously releasing vesicles (VAMP7 or vti1a), both of which are somewhat refractory to evoked release (Hua *et al*, 2011; Ramirez *et al*, 2012). Similarly, several endosomal SNAREs including vti1a, Stx6, and Stx12/13, also engage in SV recycling and may supply RRP (Hoopmann *et al*, 2010), and VAMP4 is directed to asynchronous SV pools (Raingo *et al*, 2012). However, whether and/or to what extent these presynaptic endosomal SNAREs are present within the recycling pool and consequently confer unique properties upon it, have not been determined.

Aside from the three functionally distinct SV pools described above, accumulating evidence suggests that even within the same presynaptic terminals, distinct types of SVs coexist. For instance, vesicular glutamate transporters (VGLUTs) and vesicular monoamine transporter 2, which are responsible for uptake of glutamate and biogenic monoamines, respectively, into SVs, are present on different populations of SVs (Onoa *et al*, 2010; Silm *et al*, 2019). In addition to their distinct exocytosis kinetics at a fixed stimulation frequency, exocytotic kinetics of respective SVs during repetitive stimulation appeared to differ depending on stimulation frequencies (Silm *et al*, 2019), suggesting the existence of various populations of SV recycling pools with distinct release probabilities. Although SV biogenesis mediated by adaptor protein (AP)–3 is a key step in formation of functionally distinct SV pools (Silm *et al*, 2019), it remains unknown whether differences in their molecular compositions are due to minor SV proteins, such as presynaptic endosomal SNAREs.

To gain deeper insights into the contribution of endosomal SNAREs in SV recycling, we monitored behaviors of presynaptic endosomal SNAREs conjugated C-terminally with pHluorin (also referred to as SEP)(Miesenböck *et al*, 1998; Sankaranarayanan *et al*, 2000) in a comprehensive manner in response to various stimulation protocols. We found that, among endosomal SNAREs, Stx7 is exclusively sorted to subpopulation of the SV recycling pool which responds preferentially to high–frequency repetitive stimulation. Rapid recruitment of this pool is driven by high Ca^2+^ concentrations and requires actin dynamics, resembling a previously reported SV population that rapidly replenishes RRP after synaptic depression (Alabi & Tsien, 2012). Notably, blockade of Stx7 function by overexpressing the N–terminal domain selectively abolishes this pool. Thus, our results collectively indicate a hitherto unrecognized mechanism by which the rapidly replenishing SV recycling pool that sustains synaptic transmission during high–frequency stimulation is generated by endosomal fusion events involving Stx7 during SV recycling.

## Results

### Endosomal SNARE–SEPs localize in distinct types of presynaptic vesicle compartments

To explore recycling properties of SVs carrying presynaptic endosomal SNAREs under various stimulation protocols, we utilized a pHluorin reporter (super–ecliptic pHluorin: SEP) fused to the luminal C–terminal portion of individual SNARE proteins (Fig 1A). These include five Syntaxin (Stx) family members (Stx6, 7, 8, 12/13, and 16), all of which were highly enriched in pure SV fraction (Takamori *et al*, 2006) (Appendix Fig S1). We also included two additional endosomal SNAREs, vti1a and VAMP7, which reportedly mediate spontaneous release rather than stimulus–dependent evoked release (Hua *et al*, 2011; Ramirez *et al*, 2012). As a control for authentic SV residents, two widely used SEP constructs, SypHy (Synaptophysin fused with SEP) (Fig 1A) and VGLUT1–SEP were used (Granseth *et al*, 2006; Voglmaier *et al*, 2006). The SEP reporters have been widely used to monitor SV recycling, owing to their pH dependence by which SEP fluorescence is quenched in an acidic lumen of SVs, and de–quenched upon exposure to an extracellular neutral pH by exocytosis (Figs 1B-D). When SypHy was lentivirally transduced into cultured hippocampal neurons (Egashira *et al*, 2015, 2016), it was sorted to presynaptic compartments, evidenced by immunostaining with anti–Syb2 antibody (Fig 1B). When neurons were electrically stimulated repetitively at 10 or 40 Hz (300 or 200 action potentials (APs)), SypHy fluorescence robustly increased in response to the stimuli, and > 30–40% of all SypHy molecules, estimated by application of 50 mM NH_4_Cl at the end of the recordings, was engaged in exocytosis by the end of stimulation (Figs 1D–F). On the other hand, although presynaptic endosomal SNARE–SEPs were similarly co–localized with Syb2 (Fig 1e), the vast majority of endosomal SNARE-SEPs showed relatively smaller responses compared to SypHy (Figs 1E,F; < 25%) and in some cases, responses upon 40–Hz stimulation were significantly smaller than those elicited at 10 Hz (e.g. Stx8–SEP, Stx16–SEP) (Figs 1E,F). Strikingly though, Stx7–SEP exhibited unique behavior, i.e. stimulation at 10 Hz failed to elicit a clear fluorescence increase, while 40–Hz stimulation caused a robust increase up to ~40% (Figs 1E,F). Quantitative comparisons of all SEP–reporters in response to 10–Hz and 40–Hz stimulation (with 200 APs) revealed that Stx7–SEP exhibited ~8–fold fluorescence increase at 40 Hz compared to 10 Hz (Figs 1E,F). Overall, these results indicate that different presynaptic endosomal SNARE–SEPs localize to distinct membrane compartments that are capable of recycling in an activity-dependent manner, to different degrees, at presynaptic terminals.

**Figure 1.**
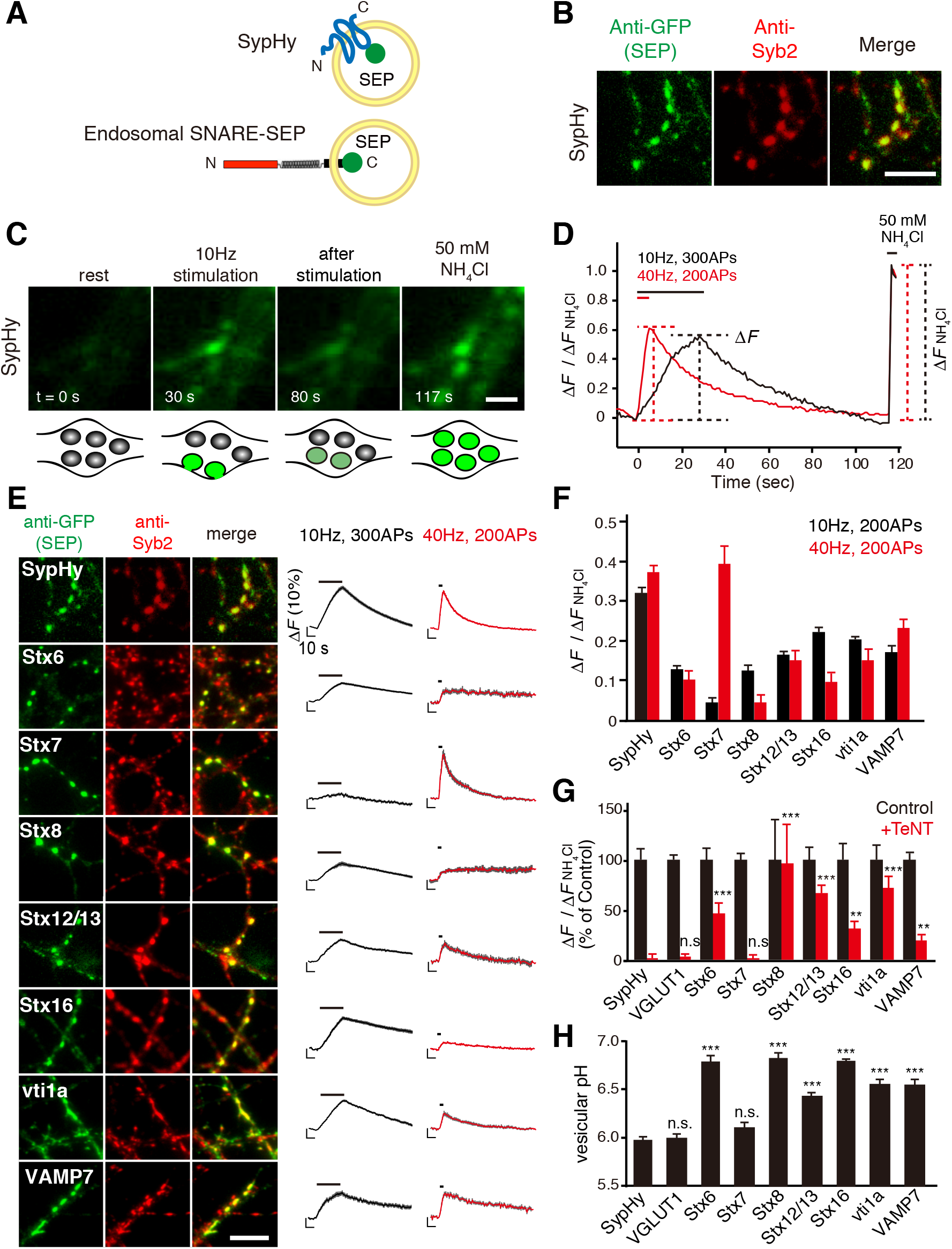
Comprehensive monitoring of presynaptic endosomal SNARE-SEPs reveals their characteristic recycling behavior. **a,** Cartoon representation of SypHy (Synaptophysin–SEP) and an endosomal–SNARE–SEP. **b–d,** Strategies for comprehensive characterization of presynaptic SNARE–SEPs. After SEP constructs were lentivirally transduced in cultured hippocampal neurons, distributions of SEP–fused proteins and their responses to repetitive electrical stimulation were analyzed by immunocytochemistry (**b**) and live fluorescence imaging (**c, d**). Representative results obtained for SypHy are shown. The SEP was visualized with anti–GFP antibody, whereas locations of presynaptic boutons were identified with anti–synaptobrevin 2 (Syb2) –antibody. To estimate fractional responses of each SEP construct, 50 mM NH_4_Cl (pH7.4) was applied at the end of recordings, which revealed total expression of SEP–fused proteins at individual boutons. Scale bars indicate 5 μm in (**b**) and 2 μm in (**c**). **e,** Synaptic localization and stimulus–dependent recycling of endosomal SNARE–SEPs upon 10–Hz or 40–Hz stimulation. Left images are representative images of each SEP–fused protein. Scale bar indicates 5 μm. Right traces show average fluorescence of individual SEP–fused proteins upon 10–Hz (300 APs; black) and 40–Hz (200 APs; red) stimulation. Fluorescence was normalized to those during NH_4_Cl application. Data are averages of > 50–200 boutons. **f,** Peak fluorescence of SEP probes at 200APs of 10–Hz (black) and 40–Hz stimulation (red). **g,** Effect of TeNT pretreatment on recycling of SEP probes in comparison to SypHy at 40–Hz stimulation (200 APs) (see also Appendix Fig S2). Data were obtained from >35–100 boutons. For bar graph comparisons, fluorescence peaks without TeNT preatment were taken as 100 %. n.s *p* > 0.05, ** *p* < 0.01, \*\*\**p* < 0.001 compared to SypHy after TeNT treatment (Student’s *t*–test). **h**, Luminal pHs of vesicle compartments carrying SEP probes calculated from experiments in Appendix Fig S3. *** *p* < 0.001 compared to that of SypHy (Student’s *t*–test).

### Only Stx7–SEP exclusively localizes to Syb2-positive SVs

Given distinct recycling behaviors of the presynaptic endosomal SNARE–SEPs compared to SypHy, we wondered to what extent these endosomal SNARE–SEPs are sorted into functionally fusion competent SVs. To test this, SEP responses were monitored after pretreatment with tetanus toxin (TeNT) that enzymatically cleaves Syb2/VAMP2 (Schiavo *et al*, 1992), thereby terminating SV exocytosis (Schoch *et al*, 2001). The robust response of Stx7–SEP at 40 Hz was completely blocked by TeNT treatment, as was observed for SypHy and VGLUT1–SEP, indicating that Stx7–SEP was sorted into Syb2/VAMP2–positive, functionally competent SVs (Fig 1G; Appendix Fig S2). By contrast, responses of other endosomal SNARE–SEPs were not completely abolished by TeNT treatment, but were only diminished to varying extents (Fig 1G, Appendix Fig S2). We also examined luminal pH of intracellular compartments bearing SEP–reporters by sequential application of an acidic solution (pH 5.5) and a 50 mM NH_4_Cl solution (Mitchell & Ryan, 2004), and subsequent electrical stimulation to ensure active synapses (Fig 1H, Appendix Fig S3). These analyses revealed that the vesicular pH of Stx7–laden vesicles (6.09 ± 0.05) was identical to those of SypHy (5.96 ± 0.04) and VGLUT1–SEP (5.98 ± 0.05), whereas those of other endosomal SNARE–SEP–laden vesicles were significantly higher (> 6.40) (Fig 1H). Thus, these results collectively demonstrate that, among presynaptic endosomal SNAREs, Stx7 is peculiar in that it exclusively localizes to a subpopulation of genuine fusion–competent SVs of the recycling pool that respond preferentially to high–frequency stimulation.

### Stx7 localizes to a small portion of SV clusters with low copy numbers at the presynaptic terminals

In order to gain further evidence for localization of Stx7 in a subpopulation of SVs, we adopted several different approaches. First, we co-stained cultured hippocampal neurons with antibodies against Stx7 and Syp, and observed them using confocal microscopy (Fig 2A). Although Stx7 immunoreactivities were apparent in cell somas and dendrites as well (Appendix Fig S4), they were also observed along axons, and partially overlapped with Syp signals (Fig 2A), indicating that intrinsic Stx7 is localized within a cluster of SVs at presynaptic terminals. To examine detailed localization of Stx7 within the terminals, we next performed immunoelectron microscopy. Since our initial trials to detect endogenous Stx7 with the same antibody did not produce reliable signals, we expressed Stx7–SEP as performed for SEP imaging, and proceeded to immunostaining using anti–se antibody. Whereas SypHy signals with the same detection method were evenly spread all over SV clusters in the terminals, Stx7–SEP signals were relatively sparse and restricted to an area within SV clusters (Fig 2B). Notably, in many instances, Stx7–SEP signals were located distant from active zone structures, which is consistent with a previous observation under STED microscopy (Wilhelm *et al*, 2014).

**Figure 2.**
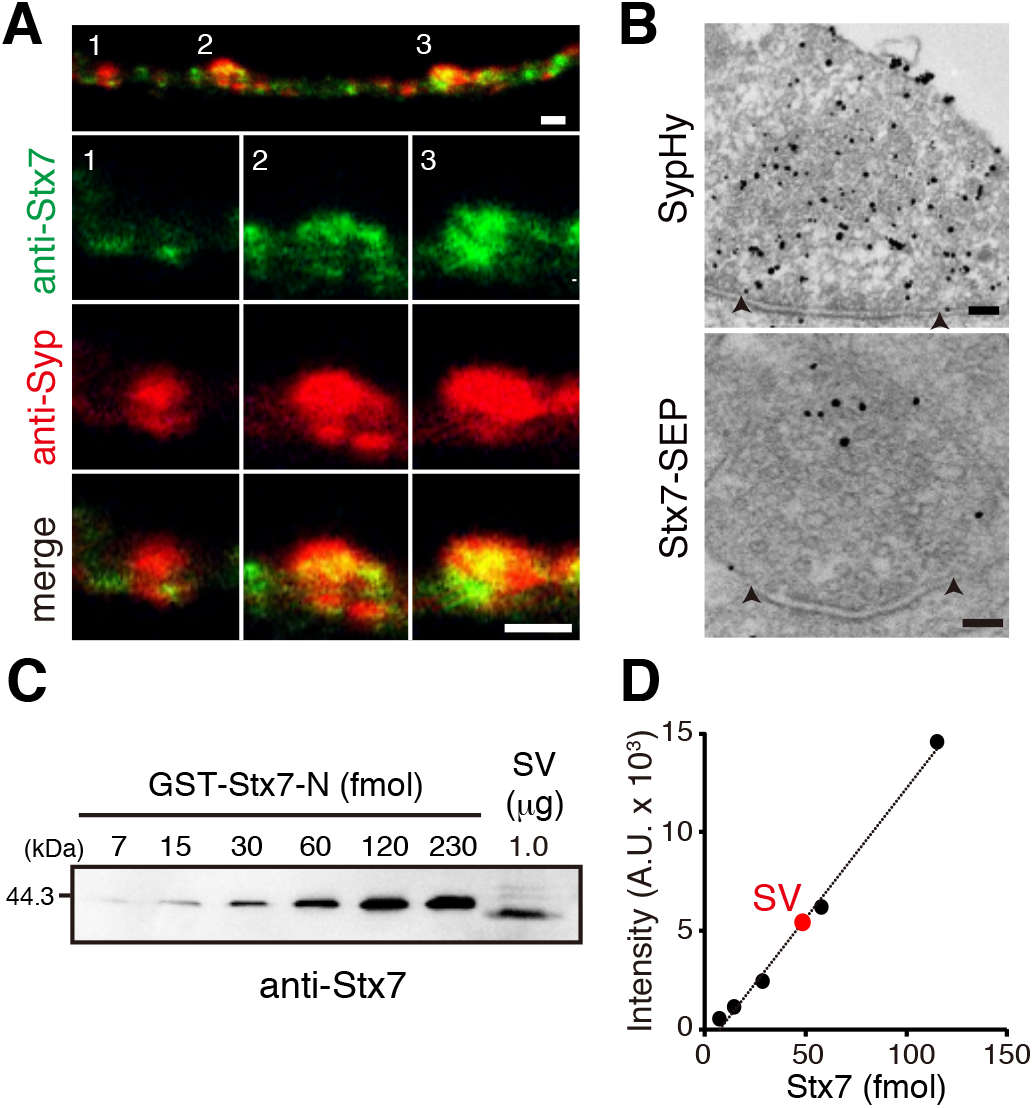
Stx7 localizes to a subpopulation of SVs at presynaptic terminals. **a,** Double immunostaining by Stx7 and Synaptophysin (Syp) antibodies in cultured hippocampal neurons at 14 DIV. An upper panel shows representative axonal localization of Stx7 (green) and Syp (red). Magnified images of numbered areas (1, 2, and 3) are shown individually below. Merged images show that Stx7 signals occupied a portion of Syp–positive puncta. Note that the specificity of the Stx7 antibody was confirmed in independent experiments where Stx7 expression level was reduced by specific shRNA (Appendix Fig. S9B). Scale bars indicate 2 μm. **b,** Electron micrographs of SypHy and Stx7–SEP at presynaptic terminals. Representative electron micrographs of a synaptic vesicle cluster labeled for SypHy (top) or Stx7–SEP (bottom) at presynaptic terminal are shown. Immunogold labeling was intensified by silver enhancement. Arrowheads indicate both ends of the active zone. Scale bars indicate 100 nm. **c,** A representative quantification of Stx7 in native SVs purified from rat brains. Various amounts of recombinant GST–Stx7–N-terminal domain (GST–Stx7–N) and a fixed amount of purified SV fraction (vesicle concentration; 26.7 nM, Protein concentration; 99.7 ng/μl) were subjected to quantitative western blot analysis (see also Appendix Fig S5 for complete datasets and control experiments for Syb2). Signal intensities of bands were quantified and plotted as a function of moles of GST–Stx7–N. Using a standard curve, copy numbers of Stx7 / SV were estimated as 0.14 ± 0.03 (SD). Data were obtained from experiments performed in quadruplicate, and the average and standard deviation are shown.

Stx7 content in the synaptosomal fraction isolated from rat brains was reported to be 78.6 copies per synapse^7^. However, the synaptosomal fraction also contains postsynaptic compartments in which Stx7 might be present (Appendix Fig S4). To estimate a fractional contribution of presynaptic Stx7 in the synaptosomal fraction, we quantified the Stx7 content in purified SVs where Stx7 was shown to be highly enriched by western blot analysis (Takamori *et al*, 2006). We estimated the copy number of Stx7 molecules per SV as 0.14 ± 0.03 (Figs 2C, D and Appendix Fig S5). Assuming that an average synaptosome contains ~380 SVs, total Stx7 molecules located on SVs is calculated to be ~53, which roughly coincides with the number of Stx7 molecules in the synaptosomal fraction (Wilhelm *et al*, 2014), suggesting that more Stx7 molecules are present in presynaptic terminals than in postsynaptic compartments.

### Stx7–SEP is sorted to SV subpopulation that requires HFS for exocytosis

One of the most striking features revealed by our comprehensive SEP imaging analysis (Figs 1E,F) was that Stx7–SEP responded preferentially to high-frequency stimulation (HFS) at 40 Hz, but rarely responded to 10–Hz stimulation. To confirm this phenomenon and to avoid any bias possibly due to heterogeneity intrinsic to each bouton or culture preparation, we continuously monitored changes in SEP fluorescence at individual boutons with various stimulation frequencies ranging from 5 Hz to 40 Hz at 5 min intervals (Fig 3A). Unlike SypHy, which reliably exhibited a robust exocytotic fluorescence increase irrespective of stimulation frequency (Fig 3A, top), Stx7–SEP hardly responded during 5–Hz or 10–Hz stimulation, while it showed robust fluorescence increases at 20 Hz and 40 Hz in the same boutons (Fig 3A, bottom).

**Figure 3.**
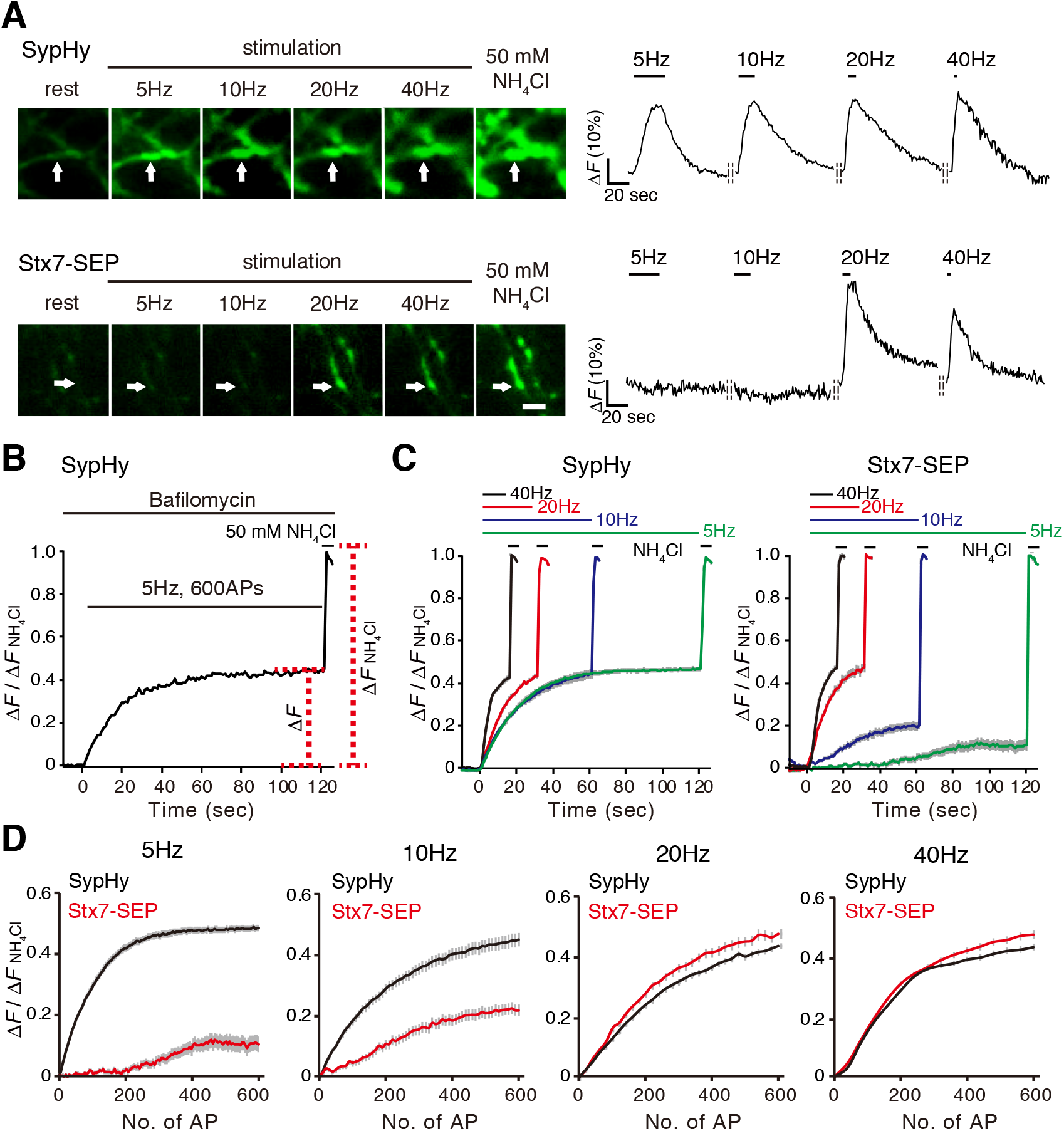
Stx7–SEP–vesicles recycle preferentially upon high–frequency stimulation. **a,** Stimulus–frequency–dependent responses of Stx7–SEP in comparison to SypHy. Neurons expressing respective SEP fusion constructs were subjected to sequential stimulation ranging from 5 to 40 Hz (200 APs) at 5 min intervals. Left images are representative images of SypHy (top), Stx7–SEP (bottom) at rest, at the end of stimulation at 5 Hz, 10 Hz, 20 Hz, and 40 Hz, and upon application of NH_4_Cl at the end of recordings. Right traces show representative traces of each SEP–fluorescence change in boutons, indicated by arrows. Data were normalized to fluorescence signals upon application of NH_4_Cl at the end of recordings. **b,** Experimental scheme to estimate kinetics of exocytosis as well as sizes of total recycling SV pools. Neurons were pretreated with 2 μM bafilomycin A1 for 60 sec and then stimulated with 600 APs at different stimulation frequencies. After cessation of stimulation, 50 mM NH_4_Cl was applied and fluorescence during NH_4_Cl application was used to normalized fluorescence signals at individual boutons. **c,** Recycling pool of SypHy and Stx7–SEP. Cells expressing SypHy (left) and Stx7–SEP (right) were pretreated with 2 μM bafilomycin A, and were stimulated with 600 APs at different frequencies (5, 10, 20 and 40 Hz). **d,** Replots of SypHy (black) and Stx7–SEP (red) responses of the results in **c** as a function of stimulus numbers (No. of AP).

We then wondered if overexpression of Stx7–SEP simply attenuated SV exocytosis generally, for instance by inactivating Ca^2+^ channels, or if Stx7–SEP localized at non-synaptic areas where non-SV type secretory vesicles that responded only to HFS were present. To exclude these possibilities, we co–expressed Stx7–SEP and Syp–mOr, in which a pH-sensitive orange fluorescent protein, mOrange2, was fused to the luminal region of synaptophysin, instead of SEP (Egashira *et al*, 2015, 2016), and monitored their fluorescence changes simultaneously in the same presynaptic boutons (Appendix Fig S6). When analyses were restricted in bouton-like structures where clear fluorescence increases of Syp-mOr were detected, Stx7-SEP rarely responded to 10–Hz stimulation (Appendix Fig S6), but exhibited drastic increase in response to 40–Hz stimulation (Appendix Fig S6).

We next wondered if more prolonged stimulation at low frequencies (5 Hz and 10 Hz) could induce exocytosis of Stx7–SEP–laden vesicles. Since endocytosis and subsequent re-acidification of SVs during prolonged stimulation might mask the SEP fluorescence increase due to SV exocytosis (Kim & Ryan, 2009), we blocked re–acidification of endocytosed vesicles with 2 μM bafilomycin A1 (Baf), a potent inhibitor of vacuolar–type H^+^–ATPases (V–ATPases), to compare exocytic properties of SypHy and Stx7–SEP (Fig 3B). We adopted 600 action potentials (APs), because that was sufficient to induce exocytosis of the entire recycling pool in cultured hippocampal neurons (Kim & Ryan, 2010). In case of SypHy, fluorescence signals plateaued at the same levels (about 40–50%) irrespective of stimulation frequencies (5–40 Hz) (Fig 3C). In stark contrast, Stx7–SEP exhibited similar exocytic properties as SypHy at 20 Hz and 40 Hz, whereas it showed much less and slower responses during 5 Hz and 10 Hz (Fig 3C). Comparisons between SypHy and Stx7–SEP by replotting as a function of stimulus numbers clearly showed that rise kinetics of Stx7–SEP were identical to those of SypHy during 20–Hz and 40–Hz stimulation, whereas those of Stx7–SEP were much slower than SypHy at 5 Hz and 10 Hz (Fig 3D).

### Stx7-vesicles represent a fraction of the recycling pool, recruitment of which requires high Ca^2+^ and actin polymerization

The fact that Stx7–SEP responded preferentially to high repetitive stimulation (> 20 Hz) suggested that exocytosis of Stx7–SEP-bearing SVs would require a high concentration of Ca^2+^. To test this hypothesis, we raised external Ca^2+^ from 2 to 8 mM, and examined whether even 10–Hz stimulation would cause robust responses. The response of SypHy was facilitated in the presence of 8 mM (Fig 4A, left), as observed previously (Chanaday & Kavalali, 2018). Notably, Stx7–SEP also exhibited a robust exocytic response to 10–Hz stimulation when external Ca^2+^ was raised to 8 mM (Fig 4A).

**Figure 4.**
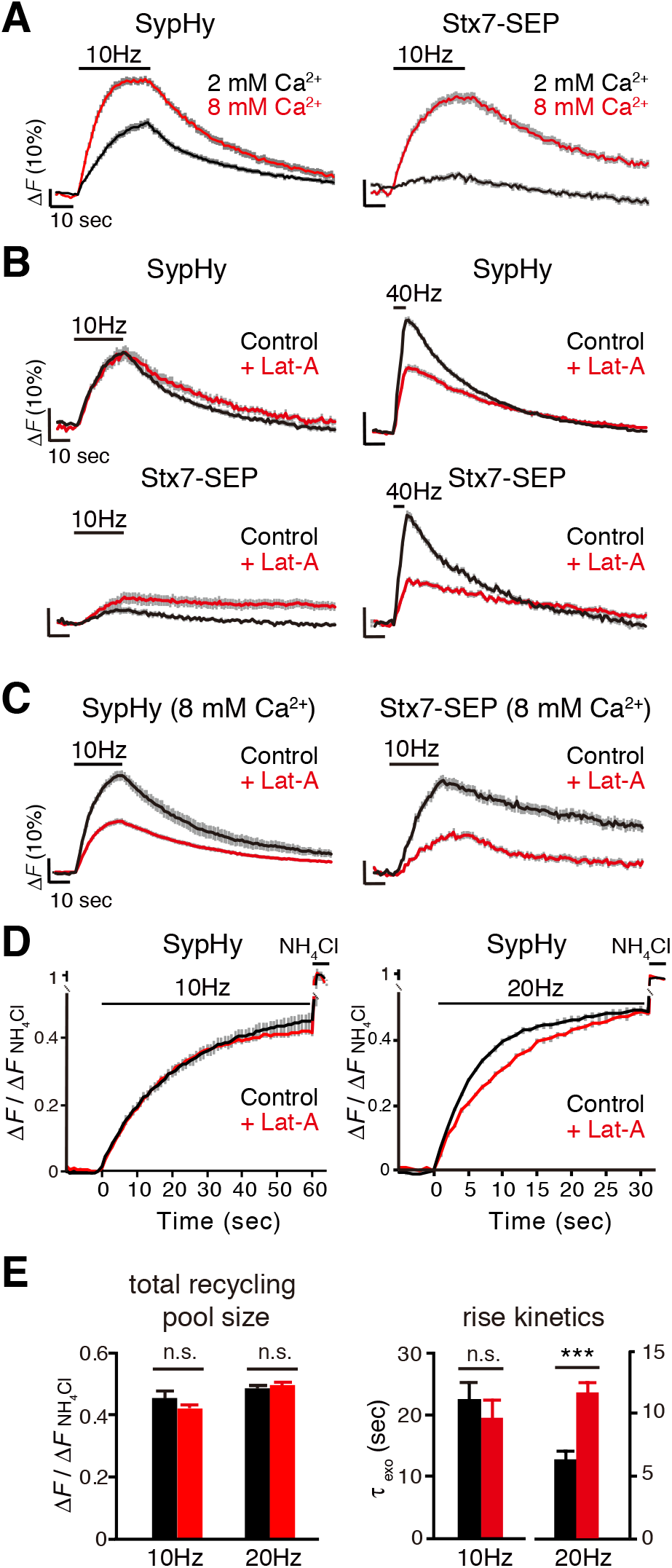
Stx7–SEP specifies a recycling pool of SVs preferentially mobilized in the presence of high Ca^2+^ concentration and actin polymerization. **a,** Responses of Stx7–SEP (right) in the presence of normal (2 mM, black) and high (8 mM, red) external Ca^2+^. For comparison, responses of SypHy under identical conditions are shown in the left panel. **b,** Effects of latrunculin A (Lat–A, 5 μM; red) on responses of SypHy (upper traces) and of Stx7–SEP (lower traces) elicited by 10–Hz (left) and 40–Hz (right) stimulation. Control experiments without Lat–A in the respective conditions are shown in black. **c,** Effects of Lat–A (red traces) on recycling of SypHy and Stx7–SEP in the presence of 8 mM external Ca^2+^. Control experiments without Lat–A in the respective conditions are shown in black. **d,** Effects of Lat–A (red) on exocytosis of total recycling pool monitored by SypHy. Responses at 10 Hz, 600APs (left) and 20 Hz, 600APs (right) with bafilomycin treatment are shown. Control experiments without Lat–A in the respective conditions are shown in black. **e**, Quantitative comparisons of total recycling pool sizes and rise kinetics in (**d)**. Normalized fluorescence peaks (left graph) or the time constants of rise time (τ_exo_) (right graph) during the 10–Hz or 20–Hz stimulation are compared. All traces are average traces from > 50–150 boutons.

We next focused on the actin cytoskeleton, because actin polymerization has often been implicated in exo–endocytosis of SVs in cultured hippocampal neurons (Wu *et al*, 2016) (but see Hua *et al*, 2011; Sankaranarayanan *et al*, 2003). In particular, Ca^2+^/calmodulin-dependent fast SV replenishment after RRP depletion is also mediated by an actin–dependent process in calyx of Held synapses (Sakaba & Neher, 2001, 2003). We therefore asked whether actin dynamics are involved in recruitment of Stx7–SEP–bearing vesicles for release during HFS. In accordance with a previous report (Hua *et al*, 2011), actin depolymerization with latrunculin A (Lat–A: 5 μM) did not show remarkable effects on SypHy recycling during 10–Hz stimulation (Fig 4B). However, Lat–A significantly retarded the SypHy response upon 40–Hz stimulation (Fig 4B, upper right), indicating that there exists a subset of SVs for which mobilization for release is actin–dependent during intense stimulation. This result is consistent with previous results reported using neurons in which actin isoforms were conditionally deleted (Wu *et al*, 2016). Notably, more pronounced reduction by Lat–A was observed for Stx7–SEP responses upon 40–Hz stimulation (Fig 4B), indicating that Stx7–SEP localizes to a subpopulation of recycling SVs for which recruitment for release is actin-dependent. Lat–B, another actin polymerization inhibitor, also decreased the SypHy and Stx7–SEP responses to similar extent at 40 Hz, but not at 10 Hz (Appendix Fig S7). Finally, when Stx7–SEP and SypHy were monitored during 10–Hz stimulation in the presence of 8 mM Ca^2+^, Lat–A significantly reduced the responses of both (Fig 4C).

To further determine whether Lat–A treatment reduced the size of the total recycling pool or whether it simply slowed SV recruitment for release, we extended stimulation lengths in the presence of Baf to measure the size of the total recycling pool of SypHy–laden vesicles (Fig 4D). As expected, kinetics measured as τ_exo_ and the total recycling pool size measured as the plateau at the end of stimulation, of SypHy responses during 10–Hz stimulation were not affected by Lat–A (Figs 4D,E), indicating that actin–dependent SV mobilization does not contribute significantly to SV mobilization at 10–Hz stimulation. In contrast, during 20–Hz stimulation, while plateaus of SypHy responses was not affected by Lat–A treatment, kinetics of SypHy responses were significantly slowed by Lat–A (Figs 4D,E).

Taken together, these results indicate that a portion of the SV recycling pool is recruited through a pathway requiring Ca^2+^–dependent actin polymerization, and that Stx7–SEP preferentially localizes to that pool.

### The N–terminal domain of Stx7 is responsible for its characteristic behavior

Like other Syntaxin family members, Stx7 comprises an N–terminal domain (NTD), the SNARE domain and a transmembrane domain (TMD) (Jahn & Scheller, 2006) (Fig 5A). To understand the molecular mechanism by which Stx7 is selectively sorted to a subpopulation of the SV recycling pool, we constructed two deletion mutants either lacking the NTD (Stx7–ΔNTD–SEP) or the SNARE motif (Stx7–ΔSNARE–SEP) (Fig 5A). When these mutants were expressed in cultured neurons, Stx7–ΔNTD–SEP was properly sorted to Syb2–positive bouton-like puncta (Fig 5B) in which it preferentially localized to the intracellular acidic compartments, with albeit higher luminal pH compared to genuine SVs (Figs 5C,D and Appendix Fig S8). However, unlike Stx7–SEP, Stx7–ΔNTD–SEP responded to 10–Hz stimulation as it did to 40–Hz stimulation (Fig 5E) and TeNT treatment did not completely abolish the responses (Fig 5F). Furthermore, responses of Stx7–ΔNTD-SEP at 40 Hz were no longer sensitive to Lat-A treatment (Fig 5G). Thus, the NTD is essential for proper sorting of Stx7 to the actin–dependent SV subpool that is preferentially recruited during HFS. By contrast, Stx7–ΔSNARE–SEP did not accumulate at Syb2–positive bouton–like puncta, and was distributed evenly on cell surfaces (Figs 5B–D and Appendix Fig S8), suggesting that the SNARE domain is indispensable for its presynaptic, as well as vesicular, localization.

**Figure 5.**
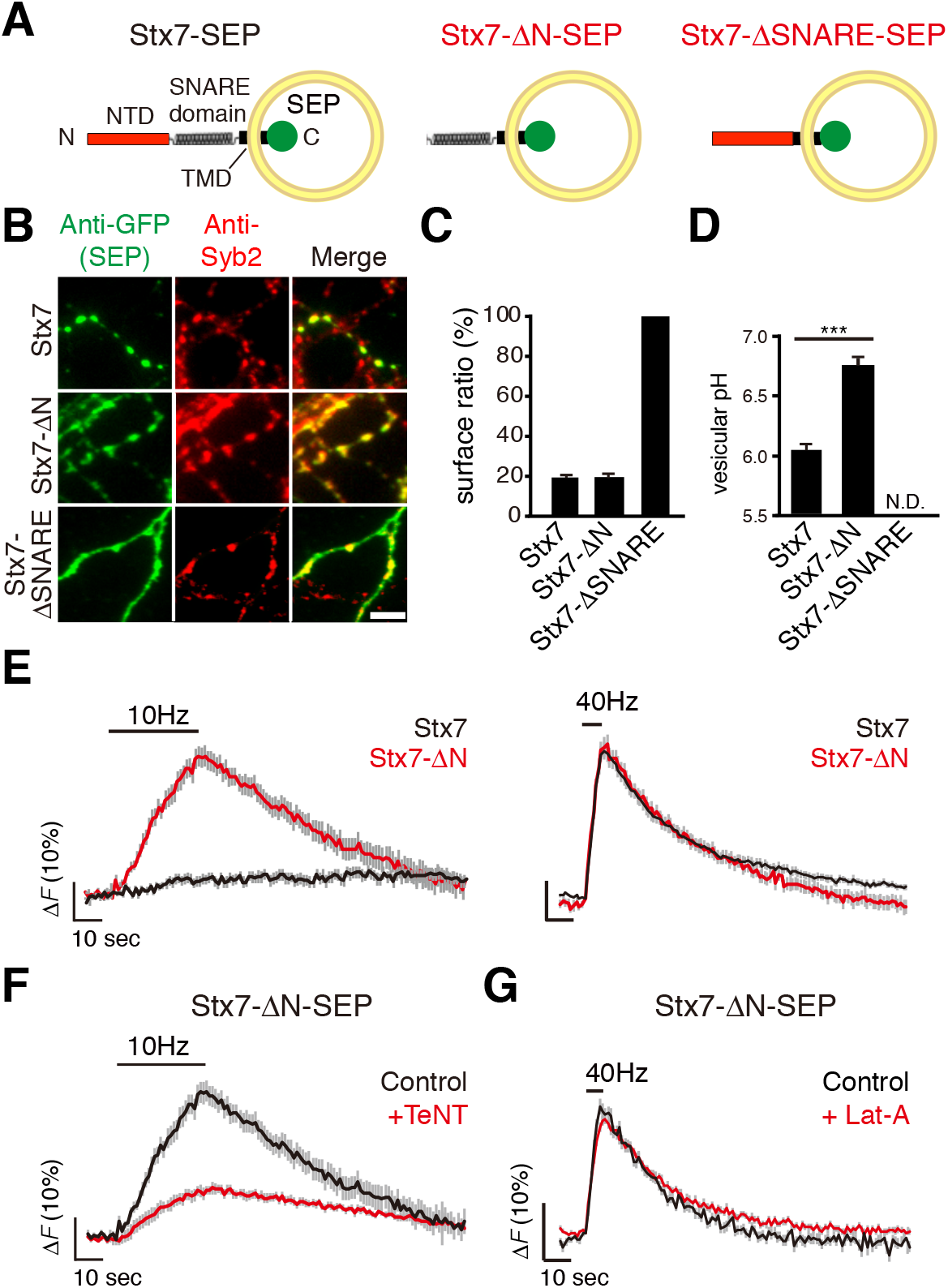
The N–terminal domain of Stx7 is required for proper sorting of Stx7 to an SV subpopulation. **a,** Schematic diagram of full length Stx7–SEP, Stx7–SEP lacking the N–terminal domain (Stx7–ΔN–SEP), and Stx7–SEP lacking SNARE motif (Stx7–ΔSNARE–SEP). **b,** Distribution of Stx7–SEP, Stx7–ΔN–SEP, and Stx7–Δ–SNARE–SEP in transfected neurons. **c,d** The surface fraction of Stx7–SEP and truncated mutants (**c**) and vesicular pHs of vesicles carrying the respective SEPs (**d**). Note that vesicular pH of Stx7–ΔSNARE–SEP could not be calculated, since it was exclusively expressed at the cell surface (N.D. indicates ‘not determined’). **e**, Responses of Stx7–SEP (black traces) and Stx7–ΔN–SEP (red traces) upon 10-Hz (left panels) and 40–Hz stimulation (right panels). **f,** Responses of Stx7–ΔN–SEP upon 10–Hz stimulation in control (black) and after TeNT treatment (red). **g**, Responses of Stx7–ΔN–SEP upon 40–Hz stimulation in control (black) and after Lat–A treatment (red).

### Overexpression of Stx7–NTD selectively reduces SV recruitment during HFS

Although results described above demonstrate that Stx7–SEP is directed to a subpopulation of the SV recycling pool for which recruitment for release requires high Ca^2+^ and actin polymerization, whether Stx7 is necessary for this subpopulation of SVs remains unclear, especially given that only a small population of SVs carries Stx7 (Fig 2). To address this question, we first examined whether silencing of Stx7 with a specific shRNA affects the SV recycling sub–pool. However, chronic knockdown of Stx7 expression severely reduced SypHy responses at 10 Hz and 40 Hz, indicating that Stx7 is necessary to establish SV pool *per se* during synapse development and maturation processes (Appendix Fig S9). As an alternative approach to inactivate Stx7, we explored specific dominant-negative effects by overexpressing the N–terminal domain of Stx7. The rationale for this is that the NTD of Stx7 interacts with its own SNARE motif, thereby inhibiting SNARE complex formation with cognate SNAREs (Antonin *et al*, 2002). In addition, the results above (Fig 5) clearly demonstrate that the NTD of Stx7 is essential for its proper sorting to the actin–dependent SV recycling pool. To this end, we placed a P2A self–cleaving peptide between the SypHy and Stx7–NTD sequences (a.a. 13–149) to ensure co-expression of both proteins (Egashira *et al*, 2016), and monitored SV recycling reported by SypHy fluorescence in transfected neurons (Fig 6A). Expression of Stx7–NTD did not alter the total recycling pool size or the kinetics of vesicle release elicited at 10 Hz (Figs 6B, C). In stark contrast, expression of the Stx7–NTD significantly slowed the increase of SypHy fluorescence during 20–Hz stimulation, while total recycling pool size was affected only slightly (Figs 6B, C). Treatment with Lat–A in addition to Stx7–NTD overexpression did not further slow release kinetics (Fig 6D), indicating direct involvement of Stx7 in the actin–dependent SV recycling pool that rapidly and preferentially replenishes RRP upon HFS.

**Figure 6.**
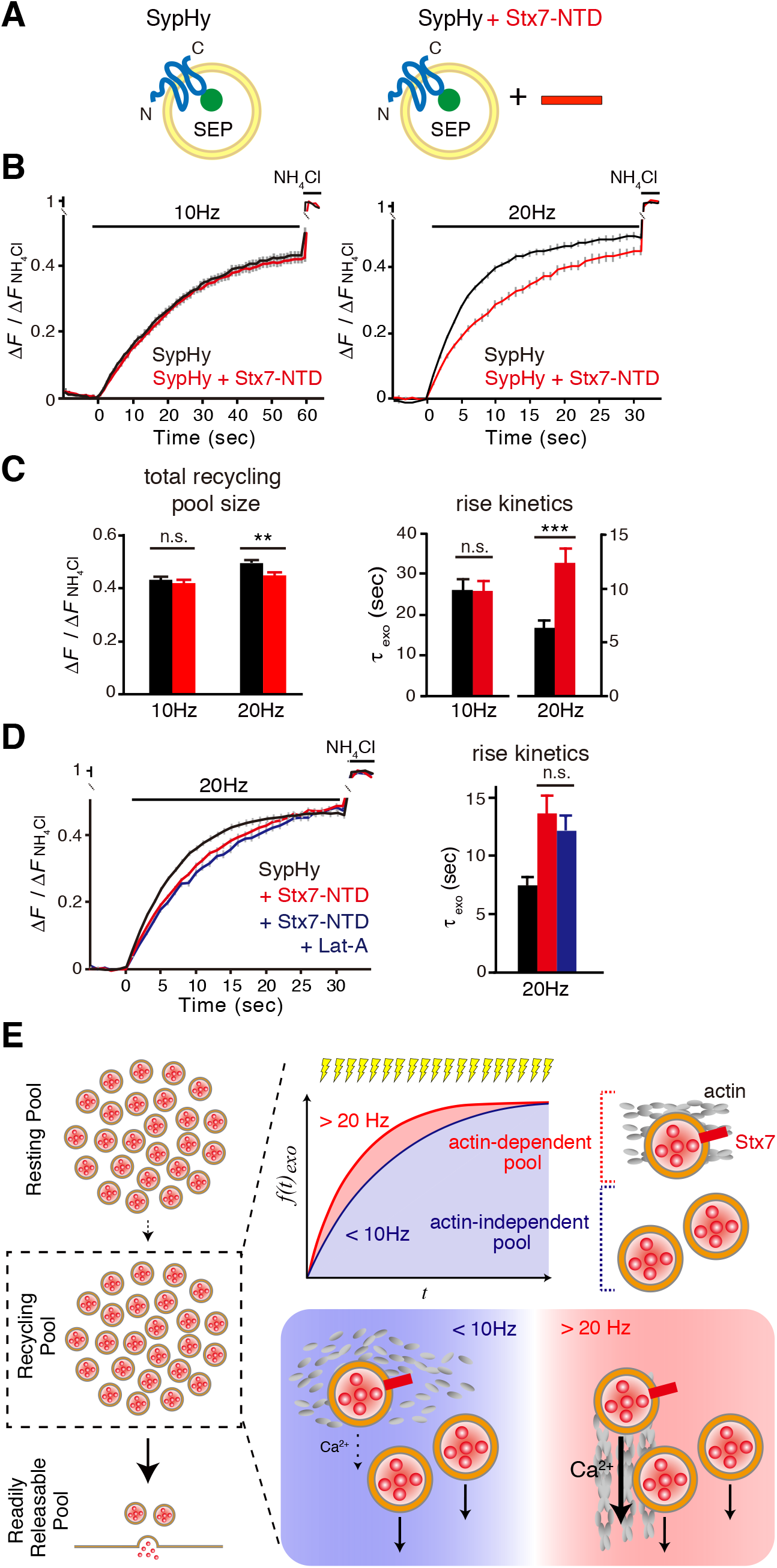
Overexpression of Stx7–NTD slowed SypHy responses only during HFS. **a,** Schematic diagram of SypHy and SypHy co–expressed with Stx7–NTD. Stx7–NTD was placed after a P2A sequence so that all SypHy–positive cells always co–expressed Stx7–NTD. **b,** Effect of Stx7–NTD on the total recycling pool monitored by SypHy responses at 10 Hz, 600APs (left) and 20 Hz, 600APs (right) after bafilomycin treatment. c, Quantification of total recycling pool sizes (left) and time constants of rise time (right, τ_exo_) in (**b**). **d,** Pretreatment with Lat–A did not further reduce release kinetics on Stx7–NTD overexpression. Responses of SypHy with Stx7–NTD in the presence of Lat–A (blue) was compared with control (SypHy only, black) and SypHy with Stx7–NTD (red) in response to at 20 Hz, 600APs. A right graph shows time constants of the rise time (τ_exo_). SypHy responses with Stx7–NTD (red) and Stx7–NTD + Lat–A (blue) did not differ significantly (*p* > 0.05).

## Discussion

Departing from comprehensive analyses of optical reporters for multiple presynaptic endosomal SNAREs, we report that an endosomal Q–SNARE, called Stx7, specifically targets a unique SV pool that can replenish RRP only during intense activity. Recruitment of this SV pool enabled by Stx7 requires high Ca^2+^ concentrations and actin polymerization, characteristics conceptually resembling a population of the recycling SV pool that rapidly replenishes RRP in calyx of Held synapses (Wu *et al*, 2016; Sakaba & Neher, 2003; Lee *et al*, 2012; Piriya Ananda Babu *et al*, 2020). Our results collectively indicate that endosomal fusion supported by Stx7, presumably together with its cognate SNAREs, is responsible for generating a functionally distinct recycling pool of SVs.

### Recycling vesicle pool heterogeneity conferred by endosomal SNAREs

Previous studies using SEP constructs have suggested that release properties of a subpopulation of SVs are dependent upon endosomal SNAREs (Hua *et al*, 2011; Ramirez *et al*, 2012; Hoopmann *et al*, 2010; Raingo *et al*, 2012). For instance, the Qc–SNARE, vti1a, and the R–SNARE, VAMP7, reside mainly on the reserve pool that is reluctantly mobilized during activity. Instead, SVs bearing these SNAREs are prone to fuse spontaneously in the absence of activity (Hua *et al*, 2011; Ramirez *et al*, 2012). Likewise, other endosomal SNAREs, Stx6, Stx12/13, and VAMP4 are capable of recycling, albeit to a lesser extent than authentic SV proteins, such as Syb2(Hoopmann *et al*, 2010; Raingo *et al*, 2012). However, our comprehensive SEP imaging analyses to monitor the effect of TeNT on stimulus-dependent fluorescence changes, as well as vesicular pHs of vesicle compartments on which they reside reveals that presynaptic endosomal SNAREs can be categorized into two distinct classes; i.e., Stx7 and other endosomal SNAREs. First, Stx7–SEP resides on a fraction of the recycling vesicle pool that is preferentially mobilized during HFS (e.g. > 20 Hz). Mobilization of this pool depends on an increase in cytosolic [Ca^2+^] exceeding a certain threshold, and also requires actin dynamics (Fig. 4). Since Stx7–laden vesicles share some characteristic features of genuine SVs, e.g., the same vesicular pH as SypHy– or VGLUT1–carrying vesicles, and complete silencing of activity–dependent exocytosis by TeNT treatment, we conclude that Stx7 resides on a subpopulation of the recycling SV pool.

In great contrast to Stx7, other presynaptic endosomal SNAREs exhibit different characteristics and behaviors compared to genuine SV residents. While they responded to wide range of stimulation, their exocytotic responses were only partially attenuated by TeNT treatment, indicating that these endosomal SNAREs are present, albeit to various extents, on Syb2/VGLUT1–negative vesicle compartments. This is further supported by the fact that the average vesicular pH of vesicles containing these endosomal SNAREs appeared to be significantly higher (~6.4) than the pH of authentic SV residents. Thus, although it was proposed that Stx6, Stx12/13, and vti1a are involved in replenishment of RRPs (Hoopmann *et al*, 2010), they are only minimally present in the RRP and recycling SV pool. Specifically, they are present in non-SV compartments, which nevertheless recycle in an activity–dependent manner. Additionally, although a previous study proposed that vti1a selectively resides on Syb2–free vesicles, which are preferentially utilized for spontaneous release rather than stimulus–dependent evoked release(Ramirez *et al*, 2012), our results reveal that the vast majority of presynaptic endosomal SNAREs are present on Syb2–free vesicles. The identity and function of such non–SV recycling vesicles at presynaptic terminals conferred by these endosomal SNAREs are enigmatic, but they may utilize non–canonical neuronal SNARE proteins and a Ca^2+^ sensor for exocytotic fusion with the plasma membrane, such as SNAP–29, or –47, Stx3 or 4, and Synaptotagmin–7 (Ibata *et al*, 2019; Jurado *et al*, 2013; Liu *et al*, 2014). Notably, recent studies have proposed the existence of non–SV–type secretory organelles at synaptic sites or in close proximity to presynaptic active zones, which secrete neuropeptides from lysosomes or dense-core vesicles (Ibata *et al*, 2019; Shimojo *et al*, 2015; Emperador-Melero *et al*, 2018). Unraveling SNAREs or Ca^2+^ sensors involved in these non–SV vesicles will enable future research to understand the physiological significance of these non–SV secretory organelles for synaptic performance and signaling.

### Actin and SV mobilization

The role of the actin cytoskeleton in regulating synaptic vesicle mobilization has long been controversial. While Lat–A, an actin polymerization inhibitor, had little impact on SV mobility or recycling in hippocampal neurons (Hua *et al*, 2011; Sankaranarayanan *et al*, 2003), similar treatment resulted in a deceleration of SV replenishment of the RRP in calyx of Held synapses (Sakaba & Neher, 2003; Lee *et al*, 2012; Piriya Ananda Babu *et al*, 2020), indicating the presence of actin–dependent rapidly replenishing SVs. Here, using SypHy imaging, we provide evidence compatible for both views regarding the role of actin dynamics in SV exocytosis and recruitment. Specifically, SVs are recruited for release in an actin–independent manner during mild stimulation, while exocytotic release of a significant fraction of SVs is accelerated by an actin–dependent process during high frequency stimulation (Figs 4B,D, and E). The former is consistent with previous observations (Hua *et al*, 2011; Sankaranarayanan *et al*, 2003) and the latter is compatible with reduced SV replenishment into the RRP proposed in calyx of Held synapses, reportedly mediated by Ca^2+^/Calmodulin-dependent processes (Sakaba & Neher, 2001). Interestingly, SVs carrying Stx7–SEP are preferentially utilized for release during intense activity and are more dependent on actin dynamics than the remainder of the recycling pool revealed by SypHy. Furthermore, recruitment of Stx7–SEP–laden SVs requires high intra–terminal Ca^2+^. Thus, our results collectively indicate that SVs characterized by Stx7 may represent a fraction of recycling SV pool that is rapidly recruited to replenish the RRP in an actin–dependent manner during high activity.

### Possible roles of Stx7 to confer the actin-dependent rapidly replenishing SVs

How does Stx7 contribute to rapidly replenishing SVs? In non–neuronal cells, Stx7 is required for homotypic/heteromeric fusion of late-endosomes and lysosomes (Mullock *et al*, 2000; Antonin *et al*, 2000; Ward *et al*, 2000) or lysosomes and autophagosomes (Matsui *et al*, 2018), and this process contributes to protein sorting and organelle maturation (He *et al*, 2016). In accordance with the role of Stx7 in non–neuronal cells, we propose that Stx7 may contribute to formation of the actin–dependent SV subpool by mediating homotypic as well as heterotypic fusion with existing endosomal structures. Recent evidence suggests that generation of clathrin–independent large endosome–like vacuoles or bulk endocytosis is the predominant form of endocytosis, especially after repetitive stimulation (Kononenko *et al*, 2014; Nicholson-Fish *et al*, 2015). Although current models predict clathrin–mediated SV reformation directly from these endosomal membranes, it is also evident that homotypic or heterotypic fusion of endosomes mediated by endosomal SNAREs can occur before SV reformation, even with multiple rounds during high activity (Holroyd *et al*, 1999; Rizzoli *et al*, 2006). Although a molecular link between Stx7 and actin is enigmatic, Stx7 may help generate the rapidly replenishing SV pool as a result of endosomal fusion and molecular sorting. This model seems to work if Stx7, but not other cognate SNAREs, are destined to SVs, and if exogenously expressed Stx7–SEP follows the trafficking route of endogenous Stx7.

Some other scenarios may explain the contribution of Stx7 to the observed results. First, it is possible that Stx7 directly or indirectly associates with actin filaments and rapid recruitment of Stx7–vesicles is only possible when the Ca^2+^/calmodulin cascade is activated. Alternatively, Stx7 may contribute to compound membrane fusion with its cognate SNAREs, involving fusion of multiple vesicles prior to a final fusion(He *et al*, 2009). However, considering the limited number of copies of Stx7 in a subpopulation of SVs (only 14% of SVs can harbor a single copy of Stx7) and the relatively high impact of Stx7–NTD overexpression on the SypHy response, as well as the highly specific behavior of Stx7 among presynaptic endosomal SNAREs, these are not likely possibilities. Further studies will be necessary to determine how Stx7, presumably with its cognate SNAREs, functionally differentiates this SV pool, thereby affecting activity–dependent synaptic performance.

## Methods

### Molecular Biology

SypHy, Syp–mOr, VGLUT1–SEP, and endosomal SNARE–SEPs were expressed in a neuron-specific manner using lentivirus–based vectors, in combination with the Tet–Off system (Egashira *et al*, 2015, 2016). Two vectors were used, a ‘regulator’ vector expressing an advanced tetracycline transactivator (tTAad) under the control of the human synapsin 1 promoter (pLenti6PW–STB), and a “response” vector that expressed SypHy, Syp–mOr, VGLUT1–SEP or endosomal SNARE–SEPs under the control of a modified tetracycline–response element (TRE) composite promoter (pLenti6PW–TRE). To construct endosomal SNARE–SEPs and deletion mutants, mouse coding sequences for Stx6 (accession no. NM_021433.3), Stx7 (accession no. NM_016797.4), Stx8 (accession no. NM_018768.2), Stx12/13 (accession no. NM_018768.2), Stx16C (accession no. NM_001102424.1), vti1a (accession no. NM_001293685.1), VAMP4 (accession no. NM_016796.3), VAMP7 (accession no. NM_001302138.1), Stx7–ΔN (a.a.168–261) and Stx7–ΔSNARE (a.a.1–167, 228–261) without the stop codons were amplified by PCR using mouse adult brain complementary DNA as a template, and fused with super–ecliptic pHluorin (SEP). Thereafter, they were cloned into a *Sma*I site located downstream of the TRE sequence in pLenti6PW–TRE using an In–Fusion Cloning Kit (Clontech)(Egashira *et al*, 2015, 2016). VGLUT1–SEP was constructed as described previously(Voglmaier *et al*, 2006) and cloned into the *Sma*I site of pLenti6PW–TRE. SypHy and Syp–mOr plasmids were generated as described previously (Egashira *et al*, 2015, 2016). To co–express SypHy and the N–terminal domain of Stx7 (Stx7–NTD; a.a.13–137), a sequence encoding a self–cleaving P2A peptide was placed between SypHy and Stx7–NTD(Egashira *et al*, 2016). To generate recombinant proteins as standards for quantification analysis, Stx7–NTD (a.a. 1–149) and Syb2–ΔTMD (a.a. 1–94) (accession no. NM_009497.3) were amplified by PCR, and cloned into the *Bam*HI site and *Sal*I site of pGEX6P–1. Lentiviral vectors carrying mCherry and either scrambled shRNA (CCTAAGGTTAAGTCGCCCTCG), mouse Stx7 shRNA–#1 (CGATATGATTGACAGCATAGA), or mouse Stx7 shRNA–#2; (GAAGCTGATATTATGGACATT) were obtained from VectorBuilder biotechnology Co. Ltd.

### Lentiviral vector production

Lentiviral vectors to express SEP constructs as well as shRNAs for Stx7 in cultured neurons were produced in human embryonic kidney (HEK) 293T cells, as described previously (Egashira *et al*, 2015, 2016). Briefly, HEK293T cells cultured in a 75–cm^2^ flask (Falcon) at ~50% confluency were transfected with 17 μg lentiviral backbone plasmids (either pLenti6PW–STB or pLenti6PW–SEP reporters) and helper plasmids (10 μg pGAG–kGP1, 5 μg pCAG–RTR2, and 5 μg pCAG–VSVG) using a calcium phosphate transfection method (Chen & Okayama, 1987). 16–24 h after transfection, culture medium was replaced with 7 mL of fresh Neurobasal medium supplemented with 2% B27 and 0.5 mM glutamine. Supernatants were collected 48 h later, and filtered through a 22–μm filter unit to remove cell debris. Aliquots were flash–frozen in liquid nitrogen and stored at −80 °C until use.

### Neuronal culture and viral expression

Primary hippocampal cultures were prepared from embryonic day 16 ICR mice, as described previously (Egashira *et al*, 2015, 2016), with slight modifications. Briefly, hippocampi were dissected, and incubated with papain (90 units/mL, Worthington) for 20 min at 37°C. After digestion, hippocampal cells were plated onto poly–D–lysine–coated coverslips framed either in 24 well plates, 12 well plates (Falcon), or a Nunc 4 well dish (Thermo fisher) at a cell density of 20,000–30,000 cells/cm^2^, and kept in a 5% CO_2_ humidified incubator. At 2–4 days *in vitro* (DIV), 40 μM FUDR (Sigma) and 100 μM uridine (Sigma) were added to inhibit the growth of glial cells. One–fifth of the culture medium was routinely replaced with fresh medium every 2–4 days until recordings. Cultures were transduced with two lentiviral vectors, the regulator vector and response vectors that encode SEP reporters or shRNAs for Stx7 at between 2 to 7 DIV, and were subjected to experiments at 12–14 DIV.

### Live imaging

Fluorescence imaging was carried out at room temperature (~27 °C) on an inverted microscope (Olympus) equipped with a 60× (1.35 NA) oil immersion objective and 75–W Xenon lamp. Cells cultured on a glass coverslip were placed in a custom–made imaging chamber on a movable stage and continuously perfused with a standard extracellular solution containing (in mM): 140 NaCl, 2.4 KCl, 10 HEPES, 10 glucose, 2 CaCl_2_, 1 MgCl_2_, 0.02 CNQX and 0.025 D–APV (pH 7.4). Field stimulation at various frequencies from 5 Hz to 40 Hz (indicated in each figure) was delivered via bipolar platinum electrodes with 1–ms constant voltage pulses (50 V). At the end of recordings, a extracellular solution containing (in mM): 50 NH_4_Cl, 90 NaCl, 2.4 KCl, 10 HEPES, 10 glucose, 2 CaCl_2_, 1 MgCl_2_ (pH 7.4) was applied directly onto the area of interest with a combination of a fast flow exchange microperfusion device and a bulb controller, both of which were controlled by Clampex 10.2. To measure kinetics of exocytosis and total recycling pool size, neurons were treated with a standard extracellular solution containing 2 μM bafilomycin A1 (Baf) (BioViotica) for 1 min, and placed in the imaging chamber. Baf–containing solution was continuously applied to neurons throughout the recordings. For HFS, we adopted a stimulation frequency of 20 Hz instead of 40 Hz, since prolonged stimulation at 40 Hz often causes unavoidable air bubbles around stimulation electrodes, which hindered reliable imaging. For TeNT experiments, neurons were incubated with 10 nM recombinant TeNT–light chain (Sigma–Aldrich) for 16 h in a CO_2_ incubator. For Lat–A and Lat–B experiments, neurons were treated with either 5 μM Lat–A (Wako) or 10 μM Lat–B (Cayman Chemical) for 1 min at RT, and subjected to imaging experiments. Lat-A or -B containing buffer was continuously supplied to neurons throughout the recordings.

To estimate luminal pH and the surface fraction of SEP probes, a low pH solution containing (in mM): 140 NaCl, 2.4 KCl, 10 MES, 4 MgCl_2_ (pH 5.5) and 50 mM NH_4_Cl solution described above were successively applied to cultured neurons as described previously(Egashira *et al*, 2015, 2016). To restrict analyses at active synaptic boutons, electrical stimulation at 40 Hz was applied for 10 s at the end of recordings.

Fluorescence images (1024 × 1024 pixels) were acquired with ORCA-Flash 4.0 sCMOS camera (Hamamatsu Photonics) in time–lapse mode either at 1 Hz (for imaging in response to stimulation) or 5.7 Hz (for estimating vesicular pH and surface fraction) under control of MetaMorph software (Molecular Devices). Fluorescence of SypHy, VGLUT1–SEP, or endosomal SNARE–SEPs was imaged with 470/22 nm excitation and 514/30 nm emission filters, and Syp–mOr was imaged with 546/556 nm excitation and 575/625 nm emission filters.

### Image data analysis

Acquired fluorescence images were analyzed using MetaMorph software. Active synapses were identified manually by changes in live measurements of fluorescence. Circular region of interests (ROIs; 2.26 μm diameter and 4 μm^2^ area) were positioned manually at the center of the highlighted fluorescence spots. Non-responsive puncta, especially for presynaptic endosomal SNARE–SEPs, which became apparent only upon NH_4_Cl application, were excluded from analysis, since they may represent endosomal SNARE-SEPs expressed outside of presynaptic terminals. To extract fluorescence changes associated with stimulation at each bouton, an average of five consecutive fluorescence values was take as F_0_, and this value was subtracted from the subsequent fluorescence values (ΔF). To estimate relative fluorescence over total SEP molecules, the peak value during NH_4_Cl application was taken as ΔF_NH4Cl_ (F_NH4Cl_ – F_0_) and used to normalize the data (ΔF/ΔF_NH4Cl_). For each constructs, fluorescence changes of at least 50 boutons were analyzed and expressed as mean ± s.e.m. To calculate rise kinetics of the fluorescence increase (τ_exo_), an average trace from one image (containing 5 – 10 active boutons) was collected, and taken as n = 1. For each averaged trace, τ_exo_ was calculated by fitting the trace with a single exponential function using a Solver function from Excel software. Averages of time constants calculated from at least 10 images were statistically evaluated.

Acquired images for vesicular pH and surface fraction measurements were also processed as described above. Vesicular pH and surface fraction were calculated as described previously (Egashira *et al*, 2015, 2016), except that fluorescence during NH_4_Cl application was approximated as fluorescence at pH 7.4. Values of p*K_a_* (7.1) and a Hill co–efficient (1.0) for SEP reported previously (Mitchell & Ryan, 2004; Sankaranarayanan *et al*, 2000) were used for calculation.

### Immunocytochemistry

Cultured neurons at DIV12–14 were fixed with 4% (wt/vol) paraformaldehyde in phosphate buffer (Wako) for 10 min at RT. After washing with PBS, cells were permeabilized with PBS containing 0.2% Triton X–100 for 20 min at RT, and incubated with PBS containing 10% (vol/vol) FBS for 30 min at RT. Cells were incubated with rabbit polyclonal anti–GFP antiserum and mouse monoclonal anti–Syb2 antibody (Cl69.1) (kind gifs from Reinhard Jahn) (Figs 1B,E, and 5B), or with rabbit polyclonal anti–Stx7 antibody (Bethyl Laboratories; A304-512A) and mouse monoclonal anti–Synaptophysin antibody (Cl7.2)(Fig 2A), or chick polyclonal anti–MAP2 antibody (1:1,000; Abcam; ab5392) and anti–Stx7 antibody (Appendix Fig S4), for 1 h at RT. Cells were rinsed 3× with PBS, and further incubated with Alexa–488–conjugated anti–rabbit IgG (1:1,500; Invitrogen), Alexa–555–conjugated anti–mouse IgG (1:1,500; Invitrogen), or Alexa–633–conjugated anti–chick IgG (1:1,500; Invitrogen) for 30 min at RT. Fluorescence images were acquired with an inverted microscope (Olympus) with an ORCA–Flash 4.0 sCMOS camera (Hamamatsu Photonics) irradiated by 75–W Xenon lamp (Alexa–488, 470/22 nm excitation and 514/30 nm emission filters; Alexa–555, 556– to 570–nm excitation and 600– to 650–nm emission filter; Alexa–633; 624– to 664–nm excitation and 692– to 732–nm emission filter) (Figs 1B,E, 5B, and Appendix Fig S4). Fluorescence images (Fig 2A) were obtained with a laser scanning confocal microscope equipped with a 100× (1.35 NA) oil immersion objective (LSM710; Zeiss), and images were collected with ZEN software.

### Immunoelectron microscopy

Hippocampal neurons grown on a coverslip were transduced with lentiviruses carrying either SypHy or Stx7–SEP and were fixed for 2 h with 4% paraformaldehyde and 0.05% glutaraldehyde in phosphate–buffered saline (PBS; pH7.4) at 15 DIV. These neurons were processed for subcellular localization with an anti–GFP antiserum by employing pre–embedding immunoelectron microscopy. Briefly, plasma membranes of fixed neurons were permeabilized with 0.1% Triton X–100 in PBS for 15 min followed by blocking with 10% fetal bovine serum in PBS for 30 min and then incubated with rabbit anti–GFP antiserum (1:1,000 dilution in 10% PBS) for 24 h at 4°C. After several washes with PBS, they were incubated with 1.4 nm immunogold–conjugated anti–rabbit IgG (Nanoprobe, 1:100 dilution in 2% normal goat serum in PBS) overnight at 4°C and then subjected to post–fixation with 1% glutaraldehyde in PBS for 10 min. Immunogold particles were enhanced for 7 min using the HQ silver enhancement kit (Nanoprobe) according to manufacturer instructions. After post–fixation with 1% osmium tetroxide in PB for 40 min on ice, these samples were stained with 1% uranyl acetate solution for 35 min, dehydrated with an ethanol series and propylene oxide for 10 min each and then embedded in Durcupan resin. After polymerization of the resin at 60 °C for 48 h, coverslips were removed and serial ultrathin sections (40 nm) were prepared. These sections were observed under a transmission electron microscope at 100 kV (H–7650, Hitachi Co., Japan) and digital images were taken with a Quemessa CCD camera (Olympus–SIS, Germany).

### Protein purification

Expression of GST–tagged Stx7–NTD or Syb2–ΔTMD was induced in *Escherichia coli* BL21 cells by adding 1 mM isopropyl–β–D–thiogalactopyranoside (IPTG) at 37°C for 4 h for GST–Stx7–NTD, or 0.1 mM IPTG at 25°C, overnight for GST–Syb2–ΔTMD. Recombinant proteins were purified with glutathione-Sepharose 4 Fast Flow (Amersham Biosciences) resin according to the manufacturer instructions. Protein concentrations were determined by BCA Assay (Pierce).

### Immunoblotting

Protein samples were separated on SDS–polyacrylamide mini–gels and blotted onto PVDF membranes (Millipore). As primary antibodies, rabbit polyclonal anti–Stx7 (Bethyl Laboratories), mouse monoclonal anti–Syb2 (Cl69.1), and mouse monoclonal anti–Tuj1 (Covance) were used. Blots were further incubated with secondary antibodies (anti–mouse IgG or anti–rabbit IgG, respectively) coupled to horseradish–peroxidase and developed using a Western Lightning Plus–ECL kit (PerkinElmer). Signals were detected using a Molecular Imager ChemiDoc (BioRad).

To estimate the copy number of Stx7 per SV, CPG–purified SV fraction (a gift from Reinhard Jahn) with known protein concentration and vesicle number, was subjected to western blot together with various amounts of standard proteins for Stx7 and Syb2. Band intensities in acquired images were quantified using Quantity One software (BioRad).

### Animals

Pregnant ICR mice were purchased from SLC, Japan. All mice were given food and water ad libitum. Animals were kept in a local animal facility with a 12–h light and 12–h dark cycle. Ambient temperature was maintained around 21 °C with a relative humidity of 50%. All animal experiments were approved by the Institutional Animal Care and Use Committee of Doshisha University.

### Statistics

All data are shown as the mean ± standard error of the mean (SEM). Unpaired *t*–tests were applied to compare means of two experimental groups. All statistical tests were two-tailed, and the level of statistical significance is indicated by asterisks: \**p* < 0.05, \*\**p* < 0.01, \*\*\**p* < 0.001. n.s.; not significant.

### Data availability

All relevant data that support the findings of this study are available from the corresponding authors upon request. The detailed data for Fig. 1E-H, 3D, E, 4, 5C-G, and 6B-D are included in Source data file.

## Acknowledgements

We thank all members of the Takamori laboratory and Dr. Tetsuya Hori for helpful discussions. We thank Steven D. Aird for editing the manuscript. This work was supported by grants from JSPS KAKENHI (16H04675, 19H03330), the JSPS Core-to-Core Program, A. Advanced Research Networks grant, and a Grant from Naito Foundation to ST, and a grant from JSPS KAKENHI (18K06473) to YM.

## Author contributions

YM and ST conceived the study. YM, KT and YF performed experiments and analyzed the data. ST wrote the manuscript with assistance from YM.

## References

Alabi AA & Tsien RW (2012) Synaptic vesicle pools and dynamics. Cold Spring Harb Perspect Biol 4: a013680

Antonin W, Dulubova I, Arac D, Pabst S, Plitzner J, Rizo J & Jahn R (2002) The N-terminal domains of syntaxin 7 and vti1b form three-helix bundles that differ in their ability to regulate SNARE complex assembly. J. Biol. Chem. 277: 36449–36456

Antonin W, Holroyd C, Fasshauer D, Pabst S, Von Mollard GF & Jahn R (2000) A SNARE complex mediating fusion of late endosomes defines conserved properties of SNARE structure and function. EMBO J. 19: 6453–6464

Chanaday NL & Kavalali ET (2018) Optical detection of three modes of endocytosis at hippocampal synapses. Elife 7:

Chen C & Okayama H (1987) High-efficiency transformation of mammalian cells by plasmid DNA. Mol. Cell. Biol. 7: 2745–2752

Egashira Y, Takase M & Takamori S (2015) Monitoring of vacuolar-type H+ ATPase-mediated proton influx into synaptic vesicles. J. Neurosci. 35: 3701–3710

Egashira Y, Takase M, Watanabe S, Ishida J, Fukamizu A, Kaneko R, Yanagawa Y & Takamori S (2016) Unique pH dynamics in GABAergic synaptic vesicles illuminates the mechanism and kinetics of GABA loading. Proc. Natl. Acad. Sci. U.S.A. 113: 10702–10707

Emperador-Melero J, Huson V, van Weering J, Bollmann C, Fischer von Mollard G, Toonen RF & Verhage M (2018) Vti1a/b regulate synaptic vesicle and dense core vesicle secretion via protein sorting at the Golgi. Nat Commun 9: 3421

Granseth B, Odermatt B, Royle SJ & Lagnado L (2006) Clathrin-mediated endocytosis is the dominant mechanism of vesicle retrieval at hippocampal synapses. Neuron 51: 773–786

He J, Johnson JL, Monfregola J, Ramadass M, Pestonjamasp K, Napolitano G, Zhang J & Catz SD (2016) Munc13-4 interacts with syntaxin 7 and regulates late endosomal maturation, endosomal signaling, and TLR9-initiated cellular responses. Mol. Biol. Cell 27: 572–587

He L, Xue L, Xu J, McNeil BD, Bai L, Melicoff E, Adachi R & Wu L-G (2009) Calcium/synaptotagmin-evoked compound fusion increases quantal size and synaptic strength. Nature 459: 93–97

Holroyd C, Kistner U, Annaert W & Jahn R (1999) Fusion of Endosomes Involved in Synaptic Vesicle Recycling. Mol Biol Cell 10: 3035–3044

Hoopmann P, Punge A, Barysch SV, Westphal V, Bückers J, Opazo F, Bethani I, Lauterbach MA, Hell SW & Rizzoli SO (2010) Endosomal sorting of readily releasable synaptic vesicles. Proc. Natl. Acad. Sci. U.S.A. 107: 19055–19060

Hua Z, Leal-Ortiz S, Foss SM, Waites CL, Garner CC, Voglmaier SM & Edwards RH (2011) v-SNARE composition distinguishes synaptic vesicle pools. Neuron 71: 474–487

Ibata K, Kono M, Narumi S, Motohashi J, Kakegawa W, Kohda K & Yuzaki M (2019) Activity-Dependent Secretion of Synaptic Organizer Cbln1 from Lysosomes in Granule Cell Axons. Neuron 102: 1184–1198.e10

Ikeda K & Bekkers JM (2009) Counting the number of releasable synaptic vesicles in a presynaptic terminal. Proc. Natl. Acad. Sci. U.S.A. 106: 2945–2950

Jahn R & Scheller RH (2006) SNAREs--engines for membrane fusion. Nat. Rev. Mol. Cell Biol. 7: 631–643

Jurado S, Goswami D, Zhang Y, Molina AJM, Südhof TC & Malenka RC (2013) LTP requires a unique postsynaptic SNARE fusion machinery. Neuron 77: 542–558

Kim SH & Ryan TA (2009) Synaptic vesicle recycling at CNS snapses without AP-2. J. Neurosci. 29: 3865–3874

Kim SH & Ryan TA (2010) CDK5 serves as a major control point in neurotransmitter release. Neuron 67: 797–809

Kononenko NL, Puchkov D, Classen GA, Walter AM, Pechstein A, Sawade L, Kaempf N, Trimbuch T, Lorenz D, Rosenmund C, Maritzen T & Haucke V (2014) Clathrin/AP-2 mediate synaptic vesicle reformation from endosome-like vacuoles but are not essential for membrane retrieval at central synapses. Neuron 82: 981–988

Lee JS, Ho W-K & Lee S-H (2012) Actin-dependent rapid recruitment of reluctant synaptic vesicles into a fast-releasing vesicle pool. Proc. Natl. Acad. Sci. U.S.A. 109: E765–774

Liu H, Bai H, Hui E, Yang L, Evans CS, Wang Z, Kwon SE & Chapman ER (2014) Synaptotagmin 7 functions as a Ca2+-sensor for synaptic vesicle replenishment. eLife 3: e01524

Matsui T, Jiang P, Nakano S, Sakamaki Y, Yamamoto H & Mizushima N (2018) Autophagosomal YKT6 is required for fusion with lysosomes independently of syntaxin 17. J. Cell Biol. 217: 2633–2645

Miesenböck G, De Angelis DA & Rothman JE (1998) Visualizing secretion and synaptic transmission with pH-sensitive green fluorescent proteins. Nature 394: 192–195

Mitchell SJ & Ryan TA (2004) Syntaxin-1A is excluded from recycling synaptic vesicles at nerve terminals. J. Neurosci. 24: 4884–4888

Mullock BM, Smith CW, Ihrke G, Bright NA, Lindsay M, Parkinson EJ, Brooks DA, Parton RG, James DE, Luzio JP & Piper RC (2000) Syntaxin 7 is localized to late endosome compartments, associates with Vamp 8, and Is required for late endosome-lysosome fusion. Mol. Biol. Cell 11: 3137–3153

Nicholson-Fish JC, Kokotos AC, Gillingwater TH, Smillie KJ & Cousin MA (2015) VAMP4 Is an Essential Cargo Molecule for Activity-Dependent Bulk Endocytosis. Neuron 88: 973–984

Onoa B, Li H, Gagnon-Bartsch JA, Elias LAB & Edwards RH (2010) Vesicular monoamine and glutamate transporters select distinct synaptic vesicle recycling pathways. J. Neurosci. 30: 7917–7927

Piriya Ananda Babu L, Wang H-Y, Eguchi K, Guillaud L & Takahashi T (2020) Microtubule and Actin Differentially Regulate Synaptic Vesicle Cycling to Maintain High-Frequency Neurotransmission. J. Neurosci. 40: 131–142

Raingo J, Khvotchev M, Liu P, Darios F, Li YC, Ramirez DMO, Adachi M, Lemieux P, Toth K, Davletov B & Kavalali ET (2012) VAMP4 directs synaptic vesicles to a pool that selectively maintains asynchronous neurotransmission. Nat. Neurosci. 15: 738–745

Ramirez DMO, Khvotchev M, Trauterman B & Kavalali ET (2012) Vti1a identifies a vesicle pool that preferentially recycles at rest and maintains spontaneous neurotransmission. Neuron 73: 121–134

Rizzoli SO, Bethani I, Zwilling D, Wenzel D, Siddiqui TJ, Brandhorst D & Jahn R (2006) Evidence for early endosome-like fusion of recently endocytosed synaptic vesicles. Traffic 7: 1163–1176

Rizzoli SO & Betz WJ (2005) Synaptic vesicle pools. Nat. Rev. Neurosci. 6: 57–69

Sakaba T & Neher E (2001) Calmodulin mediates rapid recruitment of fast-releasing synaptic vesicles at a calyx-type synapse. Neuron 32: 1119–1131

Sakaba T & Neher E (2003) Involvement of actin polymerization in vesicle recruitment at the calyx of Held synapse. J. Neurosci. 23: 837–846

Sankaranarayanan S, Atluri PP & Ryan TA (2003) Actin has a molecular scaffolding, not propulsive, role in presynaptic function. Nat. Neurosci. 6: 127–135

Sankaranarayanan S, De Angelis D, Rothman JE & Ryan TA (2000) The use of pHluorins for optical measurements of presynaptic activity. Biophys. J. 79: 2199–2208

Schiavo G, Benfenati F, Poulain B, Rossetto O, Polverino de Laureto P, DasGupta BR & Montecucco C (1992) Tetanus and botulinum-B neurotoxins block neurotransmitter release by proteolytic cleavage of synaptobrevin. Nature 359: 832–835

Schoch S, Deák F, Königstorfer A, Mozhayeva M, Sara Y, Südhof TC & Kavalali ET (2001) SNARE function analyzed in synaptobrevin/VAMP knockout mice. Science 294: 1117–1122

Shimojo M, Courchet J, Pieraut S, Torabi-Rander N, Sando R, Polleux F & Maximov A (2015) SNAREs Controlling Vesicular Release of BDNF and Development of Callosal Axons. Cell Rep 11: 1054–1066

Silm K, Yang J, Marcott PF, Asensio CS, Eriksen J, Guthrie DA, Newman AH, Ford CP & Edwards RH (2019) Synaptic Vesicle Recycling Pathway Determines Neurotransmitter Content and Release Properties. Neuron 102: 786–800.e5

Sudhof TC (2004) The synaptic vesicle cycle. Annu. Rev. Neurosci. 27: 509–547

Takamori S, Holt M, Stenius K, Lemke EA, Grønborg M, Riedel D, Urlaub H, Schenck S, Brügger B, Ringler P, Müller SA, Rammner B, Gräter F, Hub JS, De Groot BL, Mieskes G, Moriyama Y, Klingauf J, Grubmüller H, Heuser J, et al (2006) Molecular anatomy of a trafficking organelle. Cell 127: 831–846

Voglmaier SM, Kam K, Yang H, Fortin DL, Hua Z, Nicoll RA & Edwards RH (2006) Distinct endocytic pathways control the rate and extent of synaptic vesicle protein recycling. Neuron 51: 71–84

Ward DM, Pevsner J, Scullion MA, Vaughn M & Kaplan J (2000) Syntaxin 7 and VAMP-7 are soluble N-ethylmaleimide-sensitive factor attachment protein receptors required for late endosome-lysosome and homotypic lysosome fusion in alveolar macrophages. Mol. Biol. Cell 11: 2327–2333

Wilhelm BG, Mandad S, Truckenbrodt S, Kröhnert K, Schäfer C, Rammner B, Koo SJ, Claβen GA, Krauss M, Haucke V, Urlaub H & Rizzoli SO (2014) Composition of isolated synaptic boutons reveals the amounts of vesicle trafficking proteins. Science 344: 1023–1028

Wu X-S, Lee SH, Sheng J, Zhang Z, Zhao W-D, Wang D, Jin Y, Charnay P, Ervasti JM & Wu L-G (2016) Actin Is Crucial for All Kinetically Distinguishable Forms of Endocytosis at Synapses. Neuron 92: 1020–1035

